# Compensation Related Utilization of Neural Circuits Hypothesis Model in Response Inhibition in Premanifest Huntington’s Disease

**DOI:** 10.1101/495416

**Authors:** Maria V. Soloveva, Sharna D. Jamadar, Matthew Hughes, Dennis Velakoulis, Govinda Poudel, Nellie Georgiou-Karistianis

## Abstract

During premanifest stages of Huntington’s disease (pre-HD), individuals typically show increased functional brain activity thought to compensate for widespread brain anomalies. What remains unknown, is to disentangle whether increased functional brain activity reflects compensation or whether it is more of a product of HD-related pathological processes. We used a quantitative model of compensation, known as the CRUNCH (Compensation-Related Utilization of Neural Circuits Hypothesis) to characterise compensatory function in pre-HD using functional magnetic resonance imaging (fMRI). To test CRUNCH predictions, pre-HD individuals (n = 15) and age- and gender-matched controls (n = 15) performed a modified stop-signal task that incremented in four levels of stop difficulty (low, intermediate-1, intermediate-2, and high). Our results did not support the critical assumption of the CRUNCH model – controls did not show increased fMRI activity with increased level of stop difficulty. By contrast, controls showed decreased fMRI activity with increased stop difficulty in right inferior frontal gyrus pars triangularis and right caudate nucleus. Relative to controls, pre-HD individuals showed increased fMRI activity in right inferior frontal gyrus pars triangularis at intermediate-2 and high level of stop difficulty, which is the opposite effect to that predicted by the CRUNCH model. Contrary to the CRUNCH prediction, the pre-HD group showed decreased fMRI activity in right caudate nucleus at low level of stop difficulty, compared with controls. Further to this, fMRI activity patterns in right inferior frontal gyrus pars triangularis and right caudate nucleus were not accompanied by a decline in the number of successful behavioural stops in pre-HD. We therefore suggest that the CRUNCH model does not apply to characterise compensatory processes associated with response inhibition in pre-HD.

**Highlights:**

- CRUNCH does not apply to characterise compensation during stop-signal in pre-HD
- Deficits in inhibitory control in pre-HD occur 25 years prior to clinical diagnosis
- CRUNCH may show signs of selective reporting

## 1. Introduction

Huntington’s disease (HD) is an autosomal dominant progressive neurodegenerative disorder (MacDonald et al., 1993). It is commonly held that individuals with the expanded gene mutation can functionally compensate for widespread brain anomalies to support cognitive and motor function prior to clinical diagnosis (Andrews, Domínguez, Mercieca, Georgiou-Karistianis, & Stout, 2015; Gregory et al., 2017; Scheller, Minkova, Leitner, & Klöppel, 2014; Soloveva, Jamadar, Poudel, & Georgiou-Karistianis, 2018). Many functional neuroimaging studies have shown that premanifest-HD (pre-HD) individuals recruit additional brain regions and exhibit increased functional magnetic resonance imaging (fMRI) task-related activity, compared to healthy controls, during verbal working memory (Kloppel et al., 2015; Kloppel et al., 2009), spatial working memory (Georgiou-Karistianis et al., 2013a; Poudel et al., 2015), response inhibition (Rao et al., 2014), set-shifting (Gray et al., 2013), and reward processing (Malejko et al., 2014). In addition, task-related fMRI over-activation, in motor and cognitive networks, typically occurs in the context of intact behavioural performance. This pattern has been suggested to indicate a compensatory process, albeit indirectly, in the early stages of the disease during significant volumetric changes (Andrews et al., 2015; Georgiou-Karistianis, 2009; Gregory et al., 2017; Soloveva et al., 2018).

It is however difficult to conclusively argue the presence of neural compensation in pre-HD. For example, additional and/or increased fMRI activity, coupled with preserved behavioural performance, could also be a sign of: (1) neurodegeneration (e.g., Albin et al., 1992; Georgiou-Karistianis, Scahill, Tabrizi, Squitieri, & Aylward, 2013b; Rosas et al., 2008); (2) interhemispheric disinhibition (Reid, Boyd, Cunnington, & Rose, 2016); (3) medication effects (Iannetti & Wise, 2007), (4) dedifferentiation (decrease in neural specificity) (Baltes & Lindenberger, 1997); and (5) changes in neurovascular coupling affecting the physiology of the BOLD response (Hua, Unschuld, Margolis, Zijl, & Ross, 2014). As such, there have been recent recommendations for the establishment of a ‘formal’ definition of compensation in HD (i.e., Kloppel et al., 2015; Gregory et al., 2018).

Kloppel et al. (2015) defined compensation as the positive change in the relationship between fMRI activity and cognitive performance, conditional on structural disease load, as proposed by the ageing criteria of compensation (Cabeza & Dennis, 2013). They found that increased fMRI activity in right parietal cortex was positively related to better verbal working memory performance in pre-HD individuals (n = 106) close to predicted disease onset. Alternatively, for compensation to be present, Gregory et al. (2018) argued that fMRI activity and behavioural performance in HD should peak first and then gradually decrease over time with progression of brain atrophy. Compared to controls, these authors reported that pre-HD individuals and early HD patients showed a pattern of fMRI activity in dorsolateral prefrontal cortex, as well as in premotor cortex and parietal cortices, consistent with this model of compensation. While these criteria of compensation are helpful in understanding the process of functional brain re-organisation in HD, they cannot be utilised to quantitatively characterise a compensatory process in the disease.

It is established that individuals recruit additional brain regions and/or exhibit increased fMRI activity in performing a task when the brain suffers some form of structural or functional damage and when a task is cognitively demanding (Barulli & Stern, 2013; Steffener & Stern, 2012). One could argue therefore that there is a need to utilise a highly cognitively demanding task to quantitatively measure compensation in HD. More specifically, a decline in fMRI activity at higher task demands would indicate a breakdown of compensatory processes (Ji et al., 2018). Importantly, the only model of compensation proposed to date, that accounts for task difficulty in the characterisation of compensatory fMRI activity, is the Compensation-Related Utilization of Neural Circuits Hypothesis (CRUNCH), developed in the normal ageing literature (Reuter-Lorenz & Cappell, 2008).

The CRUNCH model offers a quantifiable and falsifiable test of the fMRI-behaviour relationship. We recently reviewed this model in the context of HD functional MRI literature (Soloveva et al., 2018), and suggested the CRUNCH (see Figure 1) is likely to be suited to characterise compensation in HD. Specifically, CRUNCH proposes that fMRI activity increases as task difficulty increases to compensate for task difficulty. However, when individuals reach a critical point (the ‘CRUNCH’ point), where task difficulty exceeds their resources (usually at higher task demands), fMRI activity and behavioural performance declines. In line with CRUNCH predictions, older adults exhibit increased fMRI activity at lower task demands (compensatory fMRI over-activation), compared to younger adults who instead exhibit increased fMRI activity at high task demands (Reuter-Lorenz & Cappell, 2008). Older adults reach the ‘CRUNCH’ point at intermediate levels of task demands because the compensatory mechanism breaks down, which results in decreased fMRI activity in older versus younger adults.

**Figure 1.**
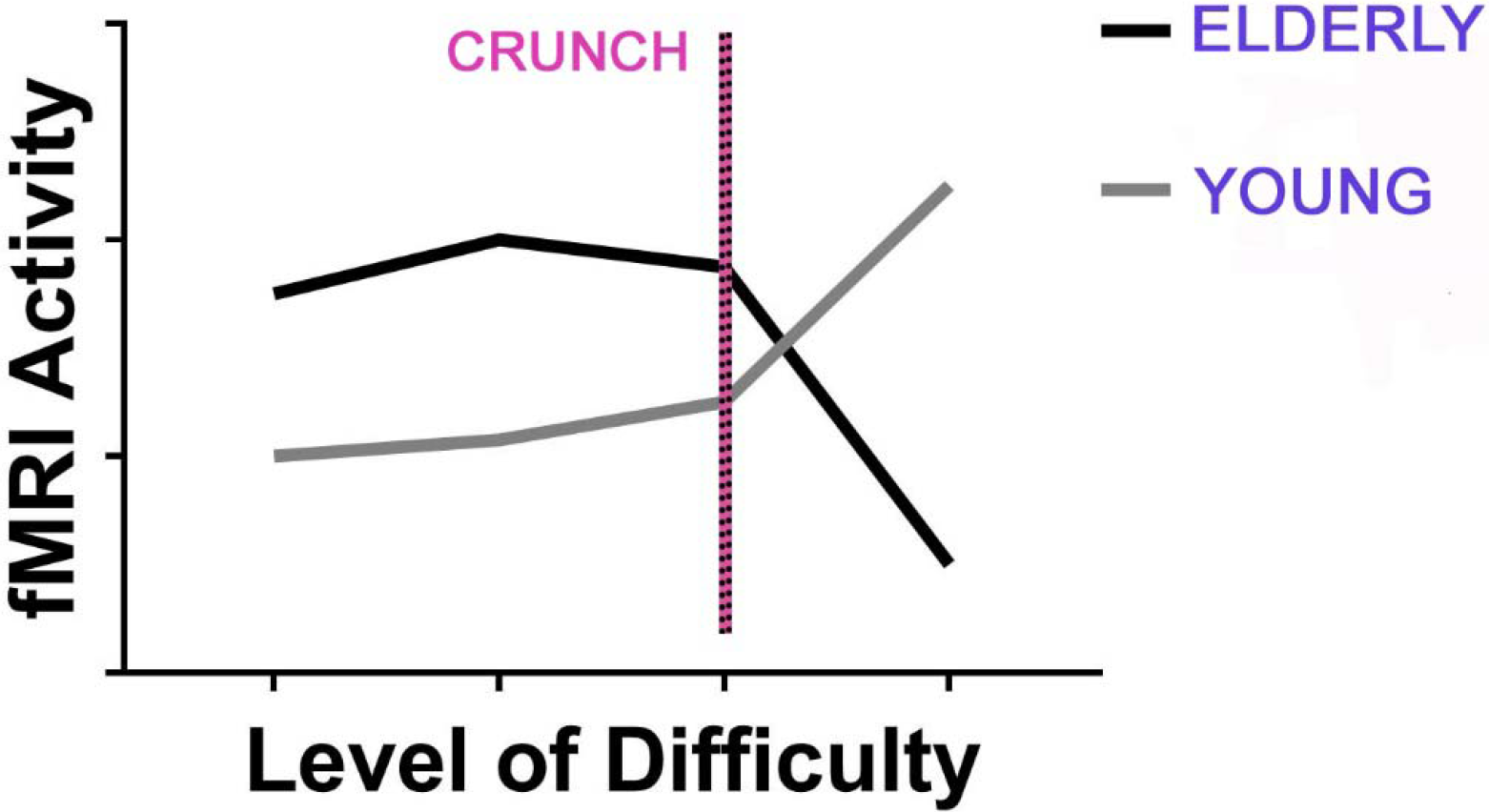
Visual representation of the hypothesised relationship between fMRI activity and task demands as proposed by the CRUNCH model (Reuter-Lorenz & Cappell, 2008). As task demands increase, older adults show increased fMRI activity. However, once task demands exceed their capacity, they reach the ‘CRUNCH’ point, and their fMRI activity and behavioural performance declines. By comparison, younger adults show increased fMRI activity at high task demands only.

Response inhibition is a hallmark of executive function (Verbruggen & Logan, 2008). In particular, the Go/No-Go task (Donders, 1969) and the stop-signal task (Logan & Cowan, 1984) are the two most commonly employed paradigms to investigate response inhibition.

In the Go/No-Go task, participants respond to a frequently presented visual ‘go’ stimulus and suppress that response to a rare ‘no-go’ stimulus. Response inhibition in the Go/No-Go task is indexed by the ‘no-go’ N2 event-related potential that relates to response conflict (Donkers & van Boxtel, 2004; Smith, Jamadar, Provost, & Mitchie, 2013; Smith, Johnstone, & Barry, 2007) and the ‘no-go’ P3 event-related potential that primarily relates to response inhibition (Smith et al., 2013; Smith et al., 2007;). During the Go/No-Go task, pre-HD individuals exhibit attenuated ‘no-go’ P3 amplitude, compared to controls (Beste, Ness, Falkenstein, & Saft, 2011; Beste, Willemssen, Saft, & Falkenstein, 2010), which is suggestive of diminished response inhibition (Smith et al., 2013; Smith et al., 2007). At the behavioural level, relative to controls, pre-HD individuals (mean of 15.66 ± 8.3 prior to predicted disease onset) also show a higher error rate on ‘no-go’ trials (Beste et al., 2011; Beste et al., 2010).

In the stop-signal task, participants respond to visual ‘go’ stimulus on ‘go’ trials as fast as possible and suppress an already initiated motor response on ‘stop’ trials following a stop signal (e.g., audio tone). To date, only a single study has examined response inhibition in HD using functional neuroimaging and the stop-signal task (Rao et al., 2014). The authors reported that pre-HD individuals close to predicted disease onset (< 7.59 years) showed reduced fMRI activity in right superior temporal gyrus and left angular gyrus, compared to controls, even though they showed no decrements in stop-signal performance. Likewise, relative to controls, pre-HD individuals far from predicted disease onset (>12.78 years) showed reduced fMRI activity in left angular gyrus/supramarginal gyrus during successful inhibition. On the other hand, unsuccessful stops were associated with decreased fMRI activity in bilateral insula and inferior frontal gyrus in the pre-HD group close to predicted disease onset; and increased fMRI activity in the right presupplementary motor area and anterior cingulate cortex in pre-HD individuals far from predicted disease onset. These results are indicative of altered functioning of brain regions that govern response inhibition in pre-HD. Given these findings, we need to clarify whether changes in the pattern of fMRI activity in inhibitory regions are a product of cortical dysfunction or indicate potential compensatory processes in pre-HD.

Evidence for impaired response inhibition is equivocal in pre-HD. In contrast to the Go/No-Go studies (e.g., Beste et al., 2011; Beste et al., 2010), where pre-HD individuals exhibited behavioural deficits in response inhibition, Rao et al. (2014) did not show decrements in stop-signal performance. It has been argued that the stop-signal task is a different index of response inhibition, compared to the Go/No-Go task. The stop-signal task measures the capacity of adults to engage in *reactive* stopping (motor response is underway (Verbruggen & Logan, 2008), whereas, in the Go/No-Go task, adults *proactively* anticipate and prepare to inhibit a response before the presentation of a ‘no-go’ stimulus. Thus, the difference in results between the Go/No-Go task and the stop-signal task in pre-HD may indicate that pre-HD individuals may be able to compensate for reactive inhibition processes but not proactive inhibition processes.

The goal of this study is to test the CRUNCH model to characterise compensatory function in pre-HD using a modified stop-signal task. In the standard stop-signal task (Logan & Cowan, 1984), participants are instructed to make fast responses on ‘go’ trials and suppress a motor response on ‘stop’ trials. We used a modified stop-signal task that parametrically increased level of difficulty, as is required to test the CRUNCH model predictions (Fabiani, 2012). The modified stop-signal task therefore comprised of low, intermediate-1, intermediate-2, and high level of difficulty. Here, to manipulate task difficulty we parametrically manipulated the perceptual load of visual stimuli on ‘go’ trials because perceptual clarity of visual information of ‘go’ stimuli can influence stopping behaviour on ‘stop’ trials (Jahfari, Ridderinkhof, & Scholte, 2013; Schmidt, Leventhal, Mallet, Chen, & Berke, 2013). Response inhibition has been modelled as the outcome of a race between independent ‘go’ and ‘stop’ processes (Verbruggen & Logan, 2009). Inhibition is successful if the ‘stop’ process finishes before the ‘go’ process, and if the ‘go’ process finishes before the ‘stop’ process, inhibition is unsuccessful (Schmidt et al., 2013). This model implies that if takes more time for an individual to process perceptually challenging ‘go’ stimuli, then it should be easier to stop the ‘go’ response after the stop-signal, compared to trials where individuals process ‘go’ stimuli quickly (not perceptually challenging). For example, using a stop-signal task, Jahfari et al. (2013) showed that making ‘go’ stimuli perceptually more challenging improved stopping performance in healthy adults.

In accordance with the CRUNCH model, we hypothesised that pre-HD individuals would exhibit increased fMRI activity at lower task demands and decreased fMRI activity at high task demands, compared with controls.

## 2. Methods

### 2.1. Participants

The study was approved by the Monash University and Melbourne Health Human Research Ethics Committees and written informed consent was obtained from each participant in accordance with the Helsinki Declaration. A total of 32 participants were recruited for the study, consisting of 17 pre-HD individuals and 15 age- and gender-matched healthy controls. We excluded two pre-HD individuals as one had head motion greater than 3mm during the MRI scan and then other reported having claustrophobia. The final sample consisted of 30 participants, including 15 pre-HD individuals (*n* = 14 right-handed; *n* = 1 left-handed) and 15 age- and gender-matched (*n* = 14 right-handed; *n* = 1 left-handed) controls. Demographic, clinical and neurocognitive data for the 30 participants are reported in Table 1.

**Table 1.** Demographic information, clinical measures and neurocognitive data.

| | Means ± SD | |
| --- | --- | --- |
|  | Controls | Pre-HD |
| n (sample sizes) | 15 | 15 |
| Gender (M:F) | 5:10 | 5:10 |
| Age | 35.60 ± 10.69 | 37.33 ± 10.82 |
| IQ estimate | 109.33 ± 6.85 | 109.07 ± 6.61 |
| UHDRS | N/A | 0.005 ± 0.02 |
| CAG repeats | N/A | 40.87 ± 2.23 |
| Disease burden score | N/A | 176.33 ± 75.88 |
| Estimated years to diagnosis | N/A | 25.77 ± 12.79 |
| SDMT | 64.47 ± 11.07 | 55.20 ± 12.60 |
| BDI-II | 3.73 ± 4.35 | 3.52 ± 3.04 |

SD, standard deviation; IQ, estimate of full-scale IQ, National Adult Reading Test (NART-2) error score; UHDRS, Unified Huntington's Disease Rating Scale (UHDRS ≤ 5 total motor score); CAG, cytosine-adenine-guanine (number of repeats $>$ 40 is full penetrance; DBS, Disease Burden Score (CAG-35.5)\*age; SDMT, Symbol Digit Modalities Test; BDI-II, Beck Depression Inventory score-II.

An independent-sample *t*-test revealed no significant differences in premorbid IQ (NART-2; Nelson, Willison, & Owen, 1992) or depression (BDI-II; Beck Steer, & Brown, 1996) scores between pre-HD individuals and controls. We did not covary for age and gender in the analysis as pre-HD individuals and controls were age- and gender-matched *a priori*. Behavioural data were analysed using SPSS Statistics v25.

A comprehensive pre-study interview was conducted with all participants to ensure that they were free from: (1) current or previous psychiatric and neurological illness (other than HD for the pre-HD group); (2) first-degree relatives with a history of severe psychiatric and neurological illness; (3) previous brain injury or stroke; (4) current or previous substance abuse disorder; (5) intellectual disability; and (6) colorblindness. Inclusion in the pre-HD group required a UHDRS total motor score of ≤ 5 (Tabrizi et al., 2009). The average pre-HD group’s estimated years to disease onset was 25.77 ± 12.79 years (Langbehn, Brinkman, Falush, Paulsen, & Hayden, 2004) and mean disease burden score (DBS) was 176.33 ± 75.88 (Orth & Schwenke, 2011). Pre-HD group reported taking serotonin–norepinephrine reuptake inhibitor (SNRI) (*n* = 1), selective serotonin and norepinephrine reuptake inhibitor (SSNRI) (*n* = 1), selective serotonin reuptake inhibitor (SSRI) (*n* = 2), and mirtazapine antidepressants (*n* = 1). One pre-HD individual was taking thyroxine to maintain normal thyroid hormone levels, as well as an atypical antipsychotic to stabilise mood and cognition and one was taking Clobazam for restless leg syndrome. Controls reported taking preventive treatment for HIV (*n* = 1) and ion supplement (*n* = 1).

All participants underwent neurocognitive and psychological testing sensitive across disease stages (Stout et al., 2011; Tabrizi et al., 2009), namely the National Adult Reading Test (NART-2; Nelson et al., 1992), the Symbol Digit Modalities Test (SDMT; Smith, 1991), and the Beck-Depression Inventory-II (BDI-II; Beck et al., 1996).

All participants performed a ~ 12-minute block of training and testing on the modified stop-signal task with low, intermediate-1, intermediate-1, and high task difficulty conditions (see Task training for details). They also completed a ~ 10-minute block of training on a visuospatial working memory task with low, intermediate-1, intermediate-2 and high difficulty conditions (results reported in Soloveva, Jamadar, Velakoulis, Poudel, & Georgiou-Karistianis, 2018). Following neurocognitive and psychological testing, as well as task training, participants underwent a 1-hour structural and functional MRI (fMRI) scan.

### 2.2. fMRI response inhibition paradigm

The fMRI experimental paradigm was a modified 22-minute stop-signal task, adapted from a standard stop-signal task (SST; see Logan & Cowan, 1984), to measure response inhibition under varied task demands between three blocks (see Figure 2 (A)). Each block started with a 500ms fixation cross followed by a series of ‘go’ and ‘stop’ trials. ‘Y’ and ‘V’ letters were used as visual stimuli on ‘go’ and ‘stop’ trials. Participants were required to make reaction time responses to ‘Y’ and ‘V’ letters (50% probability) with the right and left index finger. On 33% of trials, the stop-signal (50ms, 1000Hz audio tone) followed the ‘go’ trial after a stop signal delay (SSD) (ms) (the interval between the ‘go’ trial onset and the audio tone) and participants were required to inhibit responding when the tone occurred.

**Figure 2.**
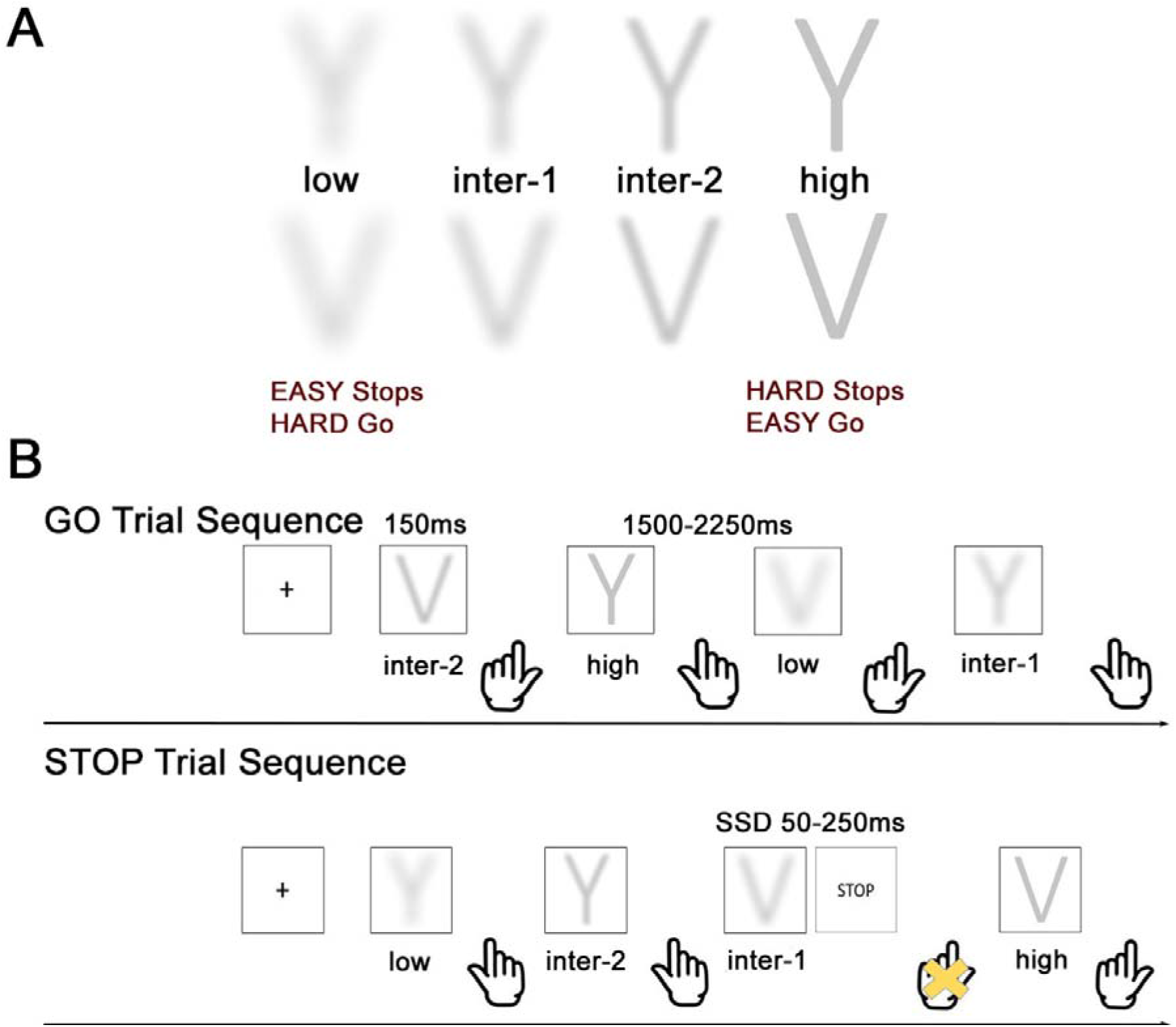
(A) ‘Y’ and ‘V’ letters subjected to high, intermediate-2, intermediate-1, and low perceptual degradation, which corresponds to low, intermediate-1, intermediate-2, and high level of stop difficulty. (B) Visual representation of the modified stop-signal task. In this example, during GO trial sequence, participants responded with the left index finger to identify the letter ‘V’ and with the right index finger to identify the letter ‘Y’ as fast and accurate as they could. During the STOP trial sequence, following a ‘go’ trial (‘V’ letter, intermediate-1 level of stop difficulty) a stop-signal audio tone randomly occurred, and the participant was required to inhibit a response. The original version of the stop-signal task was adapted from Logan and Cowan (1984).

Importantly, ‘Y’ and ‘V’ letters were subjected to four levels of perceptual degradation using Gaussian function in MATLAB 8.0 (MATLAB 8.0; http://au.mathworks.com/) (see Figure 2 (A)) to parametrically manipulate task difficulty. We labelled these conditions low, intermediate-1, intermediate-2 and high on the basis of the difficulty of the ‘stop’ process, as we were primarily interested in the response inhibition trials in this study (Figure 2A; note that the difficulty of the ‘go’ process is in the opposite direction but is not of interest in this study). Therefore, to avoid confusion, we refer to the ‘difficulty’ as the ‘stop difficulty’ manipulation. The 2-D Gaussian smoothing kernel filter with the standard deviation of 21 corresponded to low level of stop difficulty; a 2-D Gaussian smoothing kernel filter with the standard deviation of 14 corresponded to intermediate-1 level of stop difficulty; a 2-D Gaussian smoothing kernel filter with the standard deviation of 7 corresponded to intermediate-2 level of stop difficulty; and a non-smoothed image of ‘Y’ and ‘V’ letters (no 2-D Gaussian smoothing kernel filter applied) corresponded to high level of stop difficulty.

We parametrically manipulated the perceptual load of ‘Y’ and ‘V’ letters on ‘go’ trials because variance in the quality of visual information of ‘go’ stimuli will affect the ability to suppress a planned response (Jahfari et al., 2013). It is harder to make responses to a ‘go’ trial where letter identification is difficult (perceptually more challenging) vs. trials where target identification is easy (perceptually clear). As it takes more time for an individual to process a perceptually challenging ‘go’ stimulus – the ‘stop’ process finishes first, and an individual is more likely to inhibit a response on a subsequent ‘stop’ trial (Schmidt et al., 2013).

We implemented a dynamic staircase tracking procedure (Verbruggen & Logan, 2009) to individually adjust SSD based on a participant’s response to intermediate-1 level of stop difficulty. After each successful ‘stop’ trial at intermediate-1 level of stop difficulty, the SSD was extended by 50ms making it harder for a participant to inhibit on subsequent ‘stop’ trials, and after each unsuccessful ‘stop’ trial at intermediate-1 level of stop difficulty, the SSD was shortened by 50ms, making it easier for a participant to stop on subsequent ‘stop’ trials. The SSD varied from 0-250ms. The staircase tracking procedure was used in the task to ensure that participants reached ~ 50% successful inhibitions on ‘stop’ trials across all stop difficulty conditions.

Stimulus-response maps were counterbalanced across participants for the fMRI experiment. ‘Go’ and ‘stop’ trials were presented across four levels of stop difficulty (low, intermediate-1, intermediate-2, and high). Each letter was presented for 150ms and the stimulus onset asynchrony varied from 1500-2250ms with a mean of 1875ms. All responses were made via left or right button-presses using MRI-compatible response boxes. Primary outcome measures were mean RTs (ms) and the mean number of correct responses (%) on ‘go’ and ‘stop’ trials per stop difficulty condition.

### 2.3. Training task

All participants performed a ~12-minute block of training on the modified stop-signal task (see Figure 2(B)) prior to MRI scanning. This was to ensure that participants were capable of completing the task in the MRI scanner, as well as to determine the starting SSD (ms) for each participant for the MRI session. The starting SSD (ms) for each participant was calculated by averaging SSD values from intermediate-1 level of stop difficulty condition, and later used for the staircase tracking procedure during the fMRI experiment. We chose this approach to control for inter-individual variability in SSD values (Hughes, Johnston, Fulham, Budd, & Michie, 2013), so as the probability of inhibition (*P*(i)) was not influenced by SSD but rather manipulated by the perceptual load.

For the training modified stop-signal task, assignment of stimulus-response maps was counterbalanced across participants. ‘Go’ stimuli (‘Y’ and ‘V’ letters) were presented for 150ms each and the stimulus onset asynchrony varied from 1500-2250ms with a mean of 1875ms. Participants were required to press ‘Q’ and ‘]’ button presses on the keyboard with their right and left index finger to make reaction time responses to ‘Y’ and ‘V’ letters.

### 2.4. Functional data acquisition

Structural and functional MRI images were acquired using a Siemens 3-Tesla Skyra MRI scanner with a 20-channel head coil located at Monash Biomedical Imaging, Melbourne, Australia. High-resolution T1-weighted images were acquired to characterise structural integrity with sub-millimeter resolution (192 slices, 1mm slice thickness, 1mm x 1mm x 1mm voxel size, TE = 2.07ms, TR = 2300ms, FoV = 256mm; flip angle 9°). Whole-brain functional (fMRI) images were acquired during a modified stop-signal task with a gradient echo-planar sequence using interleaved slice orientation (44 slices; 3mm slice thickness, 2.5mm x 2.5mm x 3mm voxel size, TE = 30ms; TR = 2550ms; FoV = 192mm, flip angle = 80°). Time of acquisition was 1 hour per individual.

### 2.5. Data Analysis

#### 2.5.1. Behavioural data analysis

The mean RT (ms) and the mean number of correct responses (%) were computed for ‘go’ trials for each group and for each level of stop difficulty. The mean RT (ms) for ‘stop’ trials (SSRT) (ms), as well as the mean number of successful inhibitions *P*(i) (%) were estimated for each group for each stop difficulty condition separately. Group means and standard deviations for ‘go’ and ‘stop’ trials for both groups are reported in Table 2.

**Table 2.**
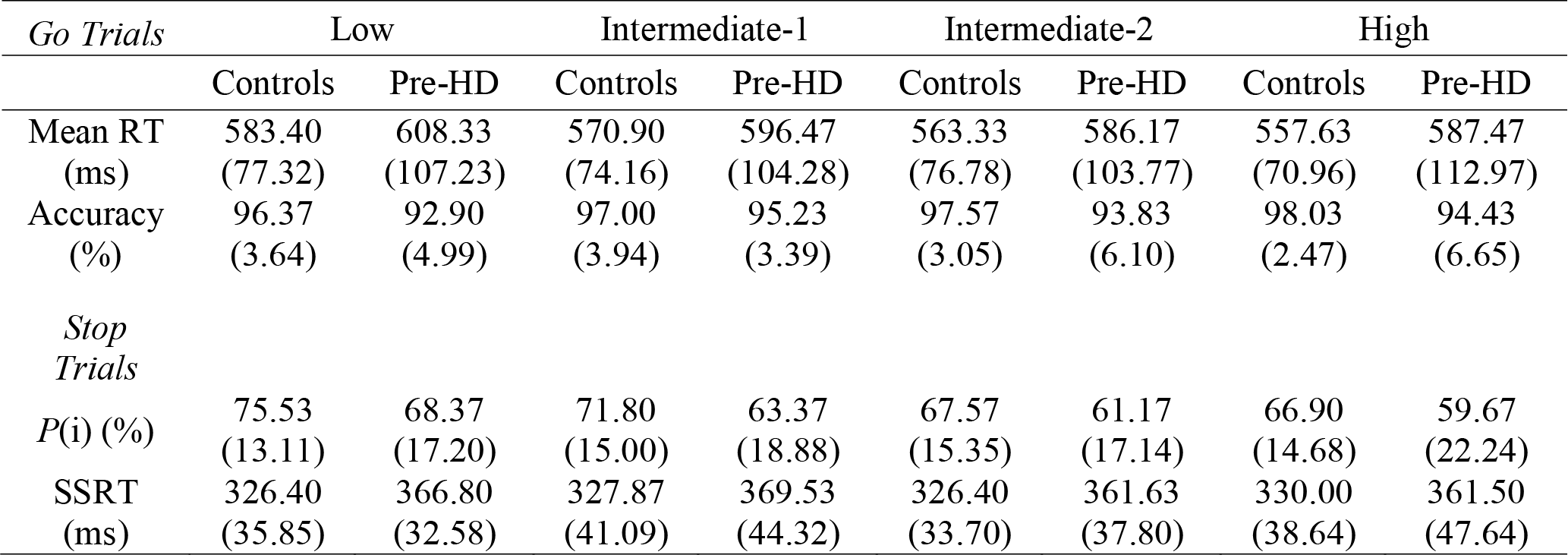
Behavioural Performance during Modified Stop-Signal Task.

We used the ‘integration method’ to calculate SSRT (ms) for each participant for each level of stop difficulty (Verbruggen & Logan, 2009). The correct ‘go’ RTs were ranked-ordered, and then the *n*th ‘go’ RT was selected, where *n* was estimated by multiplying the number of correct ‘go’ RTs in the distribution by the probability of responding to the visual stimulus (‘Y’ or ‘V’ letter) at a given SSD (ms). We used separate 2 Group (pre-HD, controls) by 4 Level of Stop Difficulty (low, intermediate-1, intermediate-2, and high) mixed model ANOVA to examine between-group differences for ‘go’ and ‘stop’ trials across each level of stop difficulty.

#### 2.5.2. fMRI pre-processing

Functional and structural MRI data were preprocessed and analysed using Statistical Parametric Mapping software (Wellcome Institute of Neurology at University College, London, UK; SPM12; http://www.fil.ion.ucl.ac.uk/spm/software/) implemented in MATLAB 8.0 (MathWorks, Natick, MA).

The first five scans of every run were removed for the magnetic field to come to equilibrium. The fMRI data underwent slice-timing correction followed by rigid-body realignment of all images to the middle slice to correct for head motion. Motion for each subject was inspected, and motion was less than 3mm for all subjects. T1-weighted structural images were segmented into separate cerebrospinal fluid, white matter, and grey matter tissue probability maps. High-resolution T1-weighted images may contain considerable amounts of non-brain tissues (e.g., eyes, skin, fat, muscle) (Fischmeister et al., 2013). We removed non-brain tissues (skull stripped) to improve the robustness of the registration process before normalisation (Fischmeister et al., 2013). The slice-time corrected and realigned fMRI scans were then co-registered to each individual’s structural T1-weighted skull-stripped image. The coregistered T1-weighted image was then spatially normalised to Montreal Neurological Institute (MNI) space using non-linear transformation routines implemented in SPM12. The resulting transformation parameters were applied to the functional images. All images were inspected visually at each step to ensure adequate data quality and processing. Finally, fMRI scans were smoothed with a 6mm full width at high maximum isotropic Gaussian kernel and high pass filtered (128s).

#### 2.5.3. Volumetric Data Analysis

We performed voxel-based morphometry (VBM; Ashburner & Friston, 2001) analysis to examine differences in brain volume between pre-HD individuals and controls. This analysis was performed to ensure that observed neural responses in each ROI varied as a function of stop difficultly rather than as a function of differences in brain volume. VBM data analyses were performed with SPM12 (Wellcome Institute of Neurology at University College, London, UK). Each participant’s unnormalised T1 structural image was segmented into grey matter (GM), white matter (WM) and cerebrospinal fluid (CSF) according to VBM protocol implemented as part of cat12 (http://www.neuro.uni-jena.de/cat/). The images were then resliced with 1.0 by 1.0 by 1.0 mm^3^ voxels producing grey and white matter images. The resulting modulated and normalised GM and WM images were further smoothed with a 6mm full width at high maximum isotropic Gaussian kernel. Modulated and normalised VBM data were used for the group comparisons of GM and WM (i.e., comparisons of an absolute amount of tissue type within a region) (Ashburner & Friston, 2001). An independent *t*-test revealed no significant differences in GM, *t*(28) = -.08, *p* = .94, WM, *t*(28) = -.66, *p* = .52, and TIV, *t*(28) = -.62, *p* = .54 between pre-HD individuals and controls, and therefore we did not include GM, WM, and TIV as regressors in the GLM.

#### 2.5.4. fMRI Analysis

Functional data were analysed within the framework of the General Linear Model (GLM) (Friston et al., 1995). Task-related fMRI signal changes were analysed for four trial types: (1) stop-correct (successful inhibition); (2) stop-incorrect (unsuccessful inhibition); (3) go-correct; and (4) go-incorrect and were included as experimental regressors of interest in GLM. On average, participants generated 121 correct ‘stop’, 58 incorrect ‘stop’, 514 correct ‘go’, and 24 incorrect ‘go’ trials. First-level models for each individual therefore comprised the four experimental regressors and six realignment parameters to control for subject motion. We computed *stop-low-correct > go-low-correct, stop-intermediate-1-correct > go-intermediate-1-correct, stop-intermediate-2-correct > go-intermediate-2-correct and stop-high-correct > go-high-correct* contrasts for each participant and subjected to a first-level analysis. The first-level analysis parameter estimates for *stop-low-correct > go-low-correct, stop-intermediate-1-correct > go-intermediate-1-correct, stop-intermediate-2-correct > go-intermediate-2-correct* and *stop-high-correct > go-high-correct* were further submitted to a second-level one-sample *t*-test to analyse fMRI activation at whole-brain level. For whole-brain level analyses, the search for fMRI activation was constrained to brain regions surviving thresholding of *p* < .001 (uncorrected) and a minimum cluster size of 50 voxels (*p* FWE corrected < .05 at cluster-level).

To examine group differences in the pattern of fMRI activity as a function of stop difficulty, we performed a region-of-interest (ROI) analysis. Parameter estimates (contrast values) were calculated for *stop-low-correct > go-low-correct, stop-intermediate-1-correct > go-intermediate-1-correct, stop-intermediate-2-correct > go-intermediate-2-correct* and *stop-high-correct > go-high-correct* contrast images. For the ROI analysis, we selected brain regions that formed part of the classic inhibitory network that showed significant fMRI activity related to inhibitory control. These ROIs were derived from a meta-analysis by Cieslik, Mueller, Eickhoff, Langner, and Eickhoff (2015) and included right and left anterior insula, right inferior frontal gyrus pars triangularis, right inferior frontal gyrus pars opercularis, right supplementary motor area, right middle frontal gyrus, right precentral gyrus, and right caudate nucleus. Further, we obtained the Montreal Neurological Institute (MNI) coordinates of the peak voxel within each ROI and extracted contrast values within 10mm sphere centred on the MNI coordinates of that peak voxel using MarsBar (Brett, Anton, Valabregue, & Poline, 2002). Finally, parameter estimates for each ROI were assessed using separate 2 Group (pre-HD, controls) by 4 Level of Stop Difficulty (low, intermediate-1, intermediate-2 and high) mixed model ANOVA (SPSS v.25).

## 3. Results

### 3.1. Behavioural results

The mean RT (ms) and the mean number of correct responses (%) on ‘go’ trials, as well as the mean RT (ms) on ‘stop’ trials (SSRT) (ms) and the mean number of successful inhibitions *P*(i) (%), for each group and for each stop difficulty condition are shown in Figure 3.

**Figure 3.**
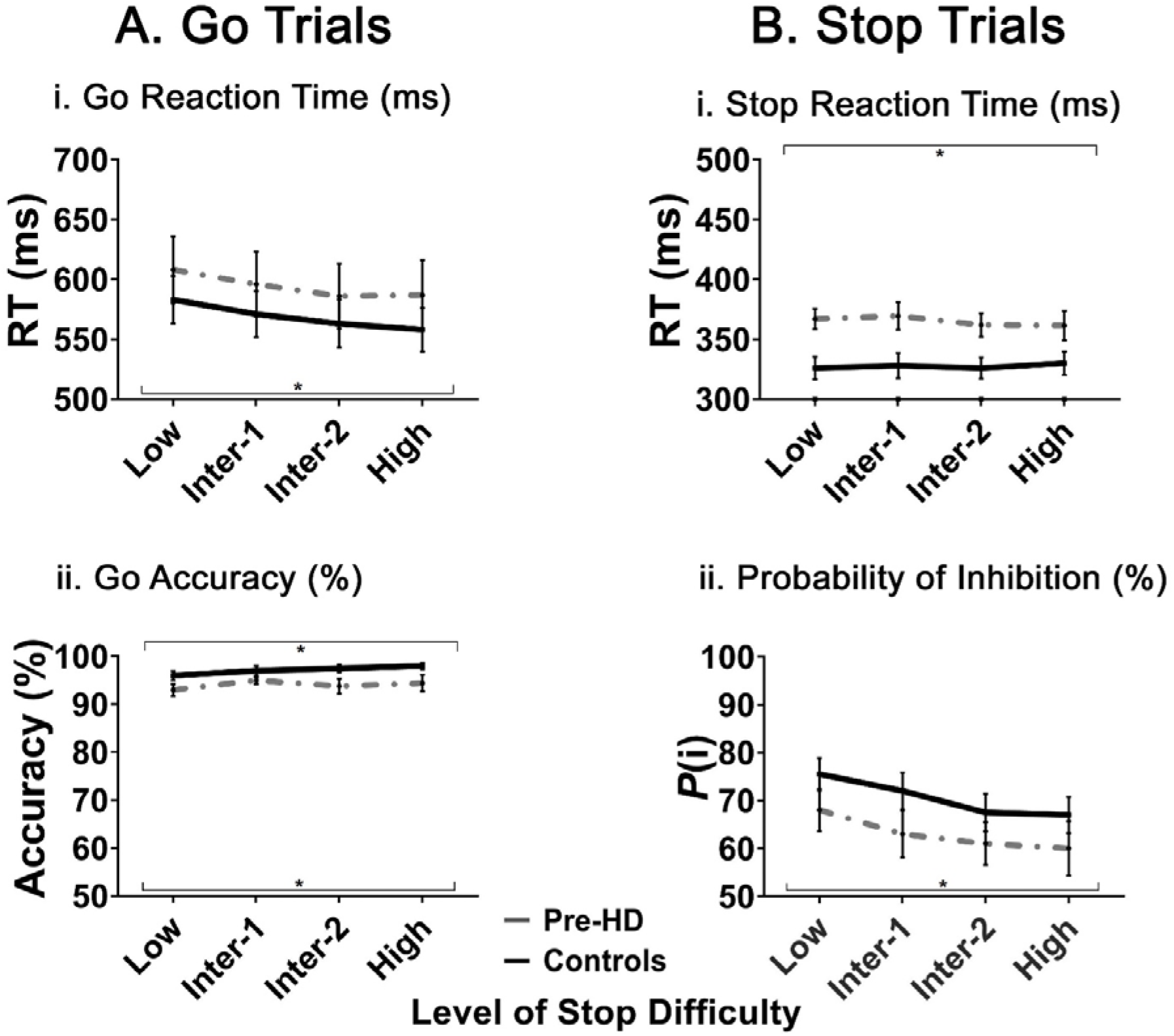
(A) Mean ‘go’ reaction times (ms) and the mean percentage (%) of correct responses on ‘go’ trials for each group for each level of stop difficulty. Standard errors are represented in the figure by the error bars for each group for each stop difficulty level. Black asterisks denote main effect of Level of Stop Difficulty on RTs for ‘go’ trials at *p* < .05 and main effect of Group on accuracy (the mean number of correct responses) (%) for ‘go’ trials at *p* < .05. (B) Mean ‘stop’ reaction times (SSRT) (ms) and the mean number of successful inhibitions *P*(i) (%) for each group for each level of stop difficulty. Standard errors are represented by the error bars for each group for each stop difficulty level. Black asterisks (*) denote main effect of Level of Group on SSRTs for ‘stop’ trials at *p* < .05 and main effect of Level of Stop Difficulty on *P*(i) for ‘stop’ trials at *p* < .05.

#### ‘Go’ Trials

Results from the mixed-design ANOVA for ‘go’ RTs revealed a significant main effect of Level of Stop Difficulty, *F*(3, 84) = 18.17, *p* < .001, indicating that both pre-HD individuals and controls were faster with increased level of stop difficulty (see Figure 3(A)). This is expected, since as Stop Difficulty increases, Go Difficulty decreases. There was no main effect of Group, *F*(1, 28) = .59, *p* = .45, confirming that RTs on ‘go’ trials were similar between the groups, and no Group by Level of Stop Difficulty interaction, *F*(3, 84) = .35, *p* = .79, indicating that RTs were not significantly different between pre-HD individuals and controls as a function of level of stop difficulty.

Results from the mixed-design ANOVA for the mean number of correct responses (%) on ‘go’ trials revealed a significant main effect of Group, *F*(1, 28) = 5.22, *p* = .03, indicating that pre-HD individuals made fewer correct responses overall, compared with controls. There was a significant main effect of Level of Stop Difficulty on the number of correct responses, *F*(3, 84) = 5.42, *p* = .002, indicating that pre-HD individuals and controls were more accurate with increased stop difficulty. This is expected, as in the ‘high’ condition the stimulus has little perceptual degradation and is easy to identify. The Group by Level of Stop Difficulty interaction was not statistically significant, *F*(3, 84) = .75, *p* = .52, suggesting that accuracy between groups did not differ as a function of stop difficulty.

#### ‘Stop’ Trials

The mixed-design ANOVA for ‘stop’ RTs demonstrated a significant main effect of Group, *F*(1, 28) = 9.00, *p* = .006, reflecting that it took longer overall for pre-HD individuals to inhibit a planned response, compared with controls (see Figure 3(B)). There was no significant main effect of Level of Stop Difficulty on SSRTs, *F*(3, 84) = .23, *p* = .88, suggesting that the time required to inhibit a response did not vary across stop difficulty conditions (see Figure (B)). The Group by Level of Stop Difficulty interaction was not significant, *F*(3, 84) = .31, *p* = .82, indicating that pre-HD individuals and controls reported similar SSRTs with increased stop difficulty.

We further found that *P*(i) decreased as a function of Level of Stop Difficulty, *F*(3, 84) = 8.73, *p* < .001, which reflects that both group were less successful in inhibiting their response when the ‘stop’ trial was perceptually clear. There was no significant main effect of Group, *F*(1, 28) = 1.62, *p* = .21, confirming that *P*(i) on ‘stop’ trials was similar between pre-HD individuals and controls. The Group by Level of Stop Difficulty interaction was not significant, *F*(3, 84) = .10, *p* = .96, suggesting that *P*(i) was not significantly different between groups with increased stop difficulty.

### 3.2. fMRI Results

#### Whole brain analysis: One-sample T-test

The whole brain analysis (*all-stop-correct > all-go-correct*) (thresholded with *p* < .001 at voxel level and *p* < .05 at FWE corrected at cluster level) averaged across groups and four levels of stop difficulty is reported in Table 3 and Figure 4. The modified stop-signal task activated the classic inhibitory network in pre-HD individuals and controls (Aron, Robbins, & Poldrack, 2014; Levy & Wagner, 2011). In particular, fMRI inhibitory-associated activations were shown throughout temporal and temporo-parietal, medial fronto-parietal and basal-ganglia-thalamo-cortical networks in both groups.

**Figure 4.**
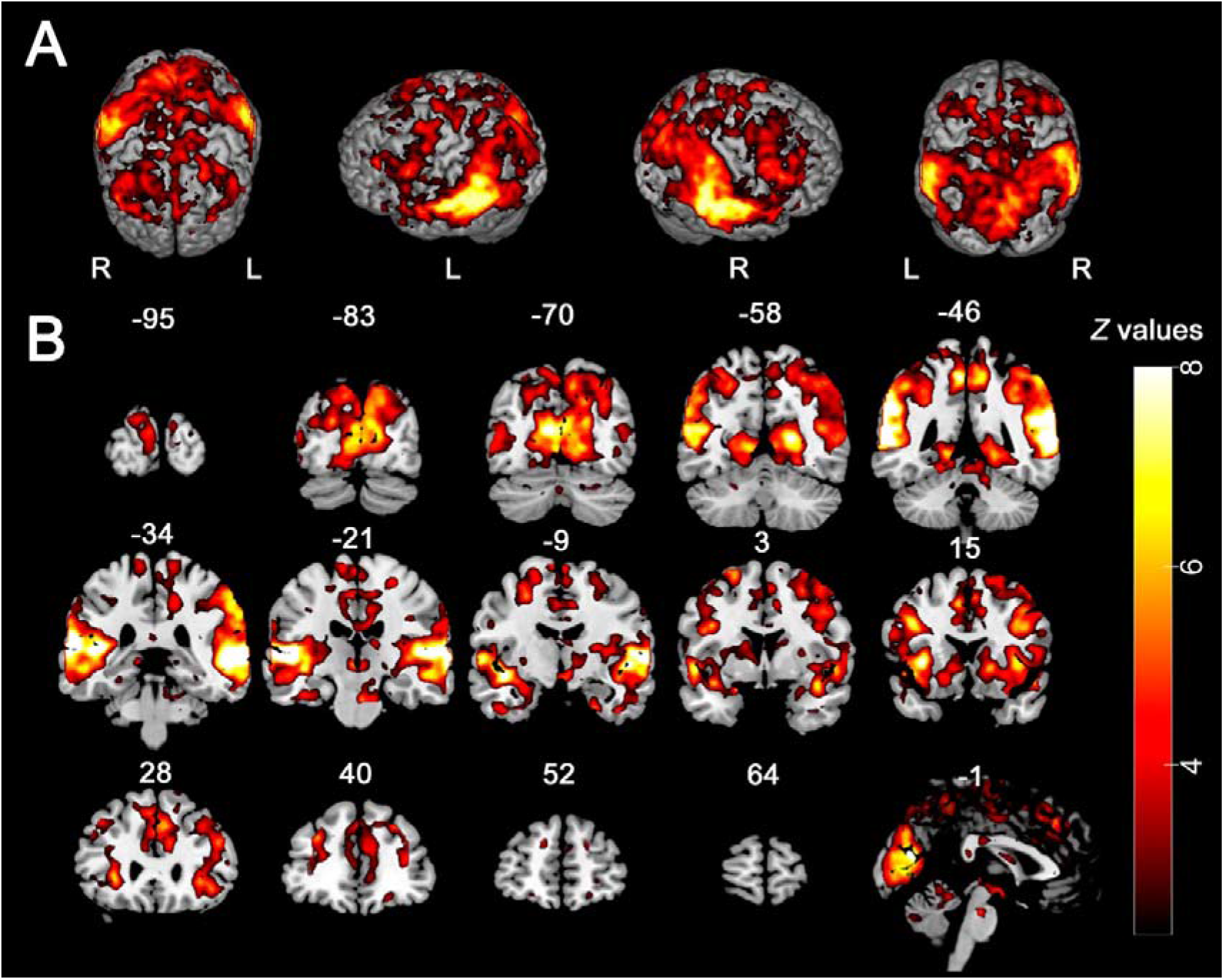
(A) Average fMRI activity across groups and conditions in contrast *all-stop-correct > all-go-correct* at *p* <.001 uncorrected, cluster level *p* <.05 FWE (whole brain analysis thresholded at *Z* ≥ 2.3, rendered view. (B) Slice view of average fMRI activity across groups and conditions in contrast *all-stop-correct > all-go-correct* at *p* <.001 uncorrected, cluster level *p* < .05 FWE (whole brain analysis thresholded at *Z* ≥ 2.3, slice view).

**Table 3.**
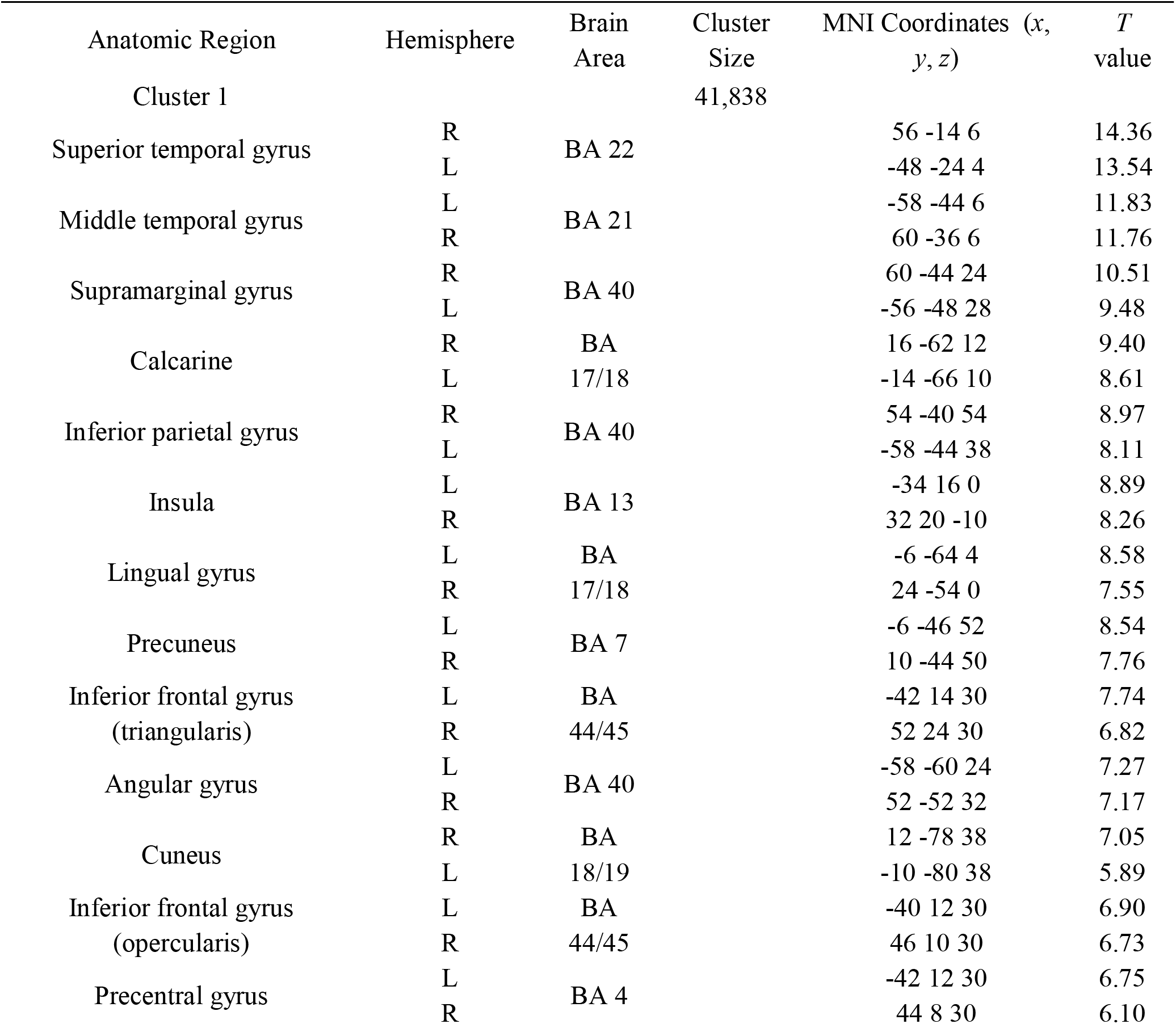
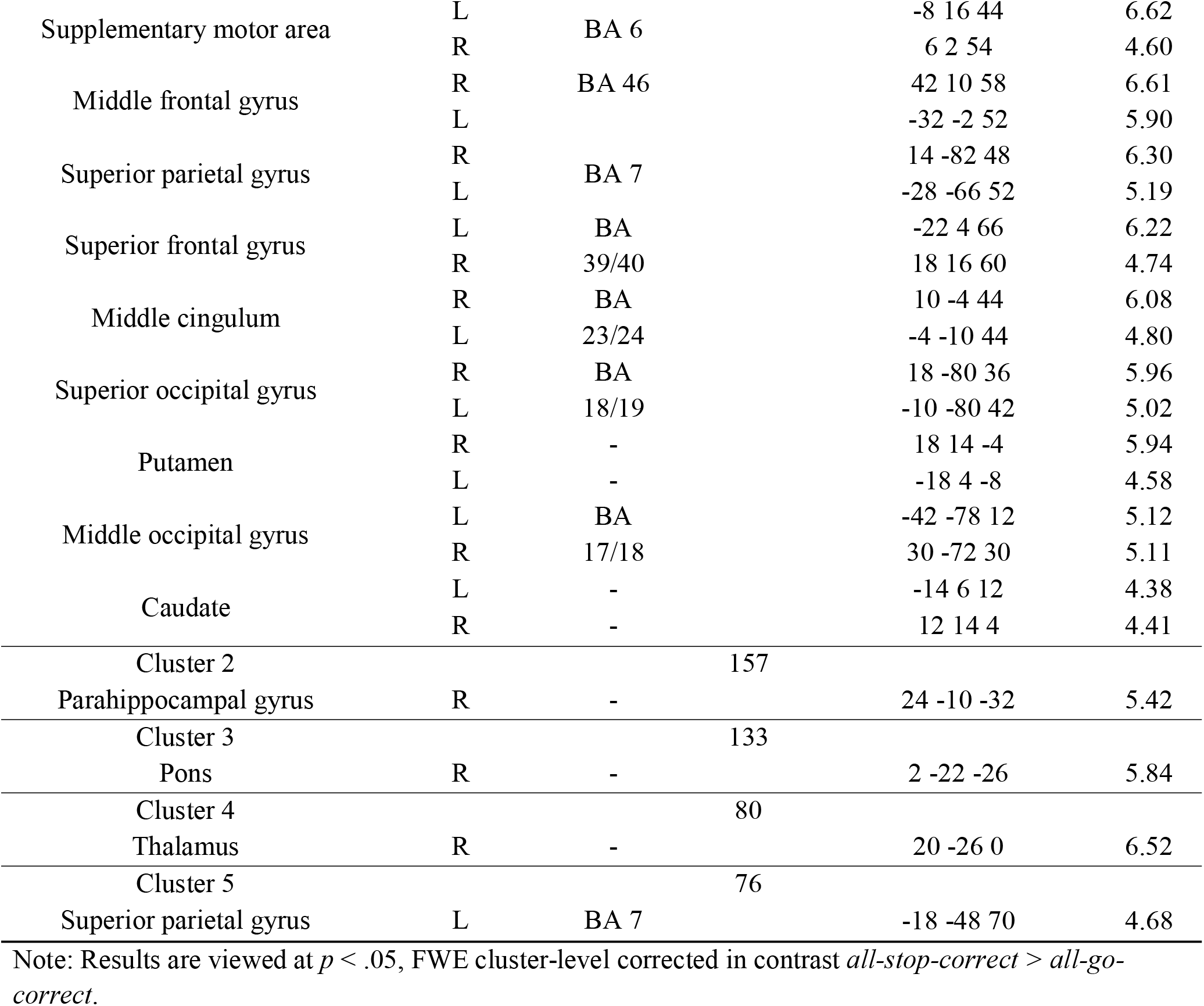
Brain regions showing signifìcant fMRI activity in pre-HD and controls in contrast all-stop-correct > all-go-correct.

#### Region of Interest Analysis

We performed 2 Group (pre-HD, controls) by 4 Level of Stop Difficulty (low, intermediate-1, intermediate-2 and high) mixed-design ANOVA to assess group differences in the pattern of fMRI activity in pre-defined ROIs, which included: (1) right and left anterior insula; (2) right inferior frontal gyrus pars triangularis; (3) right inferior frontal gyrus pars opercularis; (4) right supplementary motor area; (5) right middle frontal gyrus; (6) right precentral gyrus; and (7) right caudate nucleus. We applied a Bonferroni-correction at *p* = .00625 for testing eight comparisons.

There was no main effect of Group or Level of Stop Difficulty for any of the ROIs. The 2 Group (pre-HD, controls) by 4 Level of Stop Difficulty (low, intermediate-1, intermediate-2, and high) mixed-design ANOVA revealed a marginally significant Group by Level of Stop Difficulty interaction in right inferior frontal gyrus pars triangularis, *F*(2.68; 74.98) = 2.86, *p* = .048, *η*^2^_p_ = .09 (see Figure 5) (this did not survive a Bonferroni-correction at *p* = .00625). Simple effects analyses revealed that pre-HD individuals showed increased fMRI activity in right inferior frontal gyrus pars triangularis at intermediate-2, *F*(1, 28) = 6.45, *p* = .017, and high level of stop difficulty, *F*(1, 28) = 5.87, *p* = .02, compared with controls. However, there was no significant difference in fMRI activity between pre-HD individuals and controls in right inferior frontal gyrus pars triangularis at low and intermediate-1 level of stop difficulty, *p* = .32 and *p* = .73, respectively.

**Figure 5.**
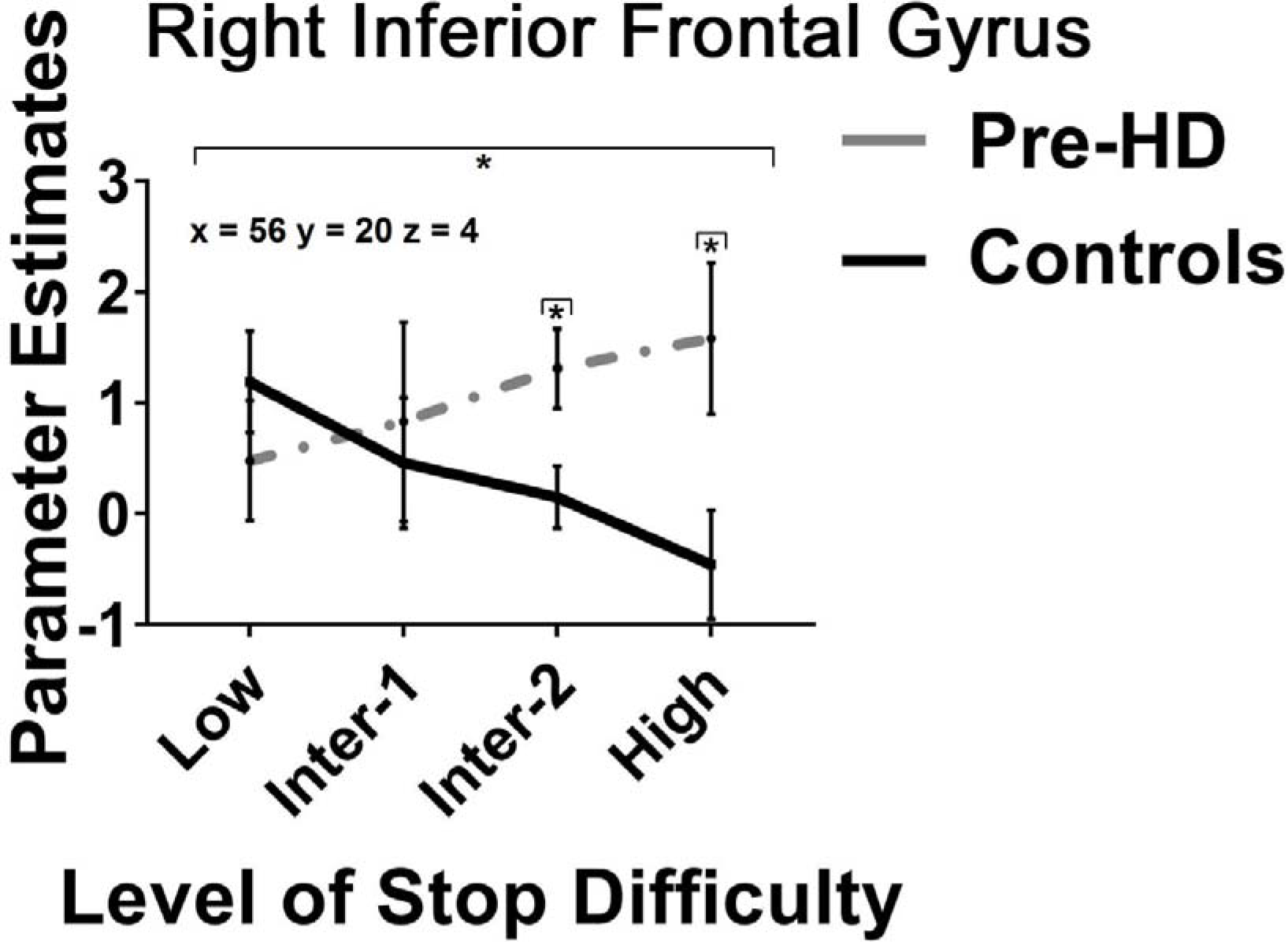
fMRI activity in right inferior frontal gyrus pars triangularis showing significant Group by Level of Stop Difficulty interaction, suggesting that the pattern of fMRI activity in this brain region varied between pre-HD individuals and controls as a function of stop difficulty. Standard errors are represented by error bars for each group and for each level of stop difficulty. Black asterisks (*) denote a significant Group by Level of Stop Difficulty interaction at *p* < .05.

The within-subjects trend analysis revealed a marginally significant linear trend for the Group by Level of Stop Difficulty interaction in right caudate nucleus, *F*(1, 28) = 4.62, *p* = .04, *η*^2^_p_ = .14 (see Figure 6) (this did not survive a Bonferroni-correction at *p* = .00625). Simple effects analyses revealed that, relative to controls, pre-HD individuals showed decreased fMRI activity in right caudate nucleus at low level of stop difficulty, *F*(1, 28) = 4.93, *p* = .04. There was no significant difference in the pattern of fMRI activity in right caudate nucleus between the pre-HD group and controls at intermediate-1, *p* = .93, intermediate-2, *p* = .12, and high level of stop difficulty, *p* = .31.

**Figure 6.**
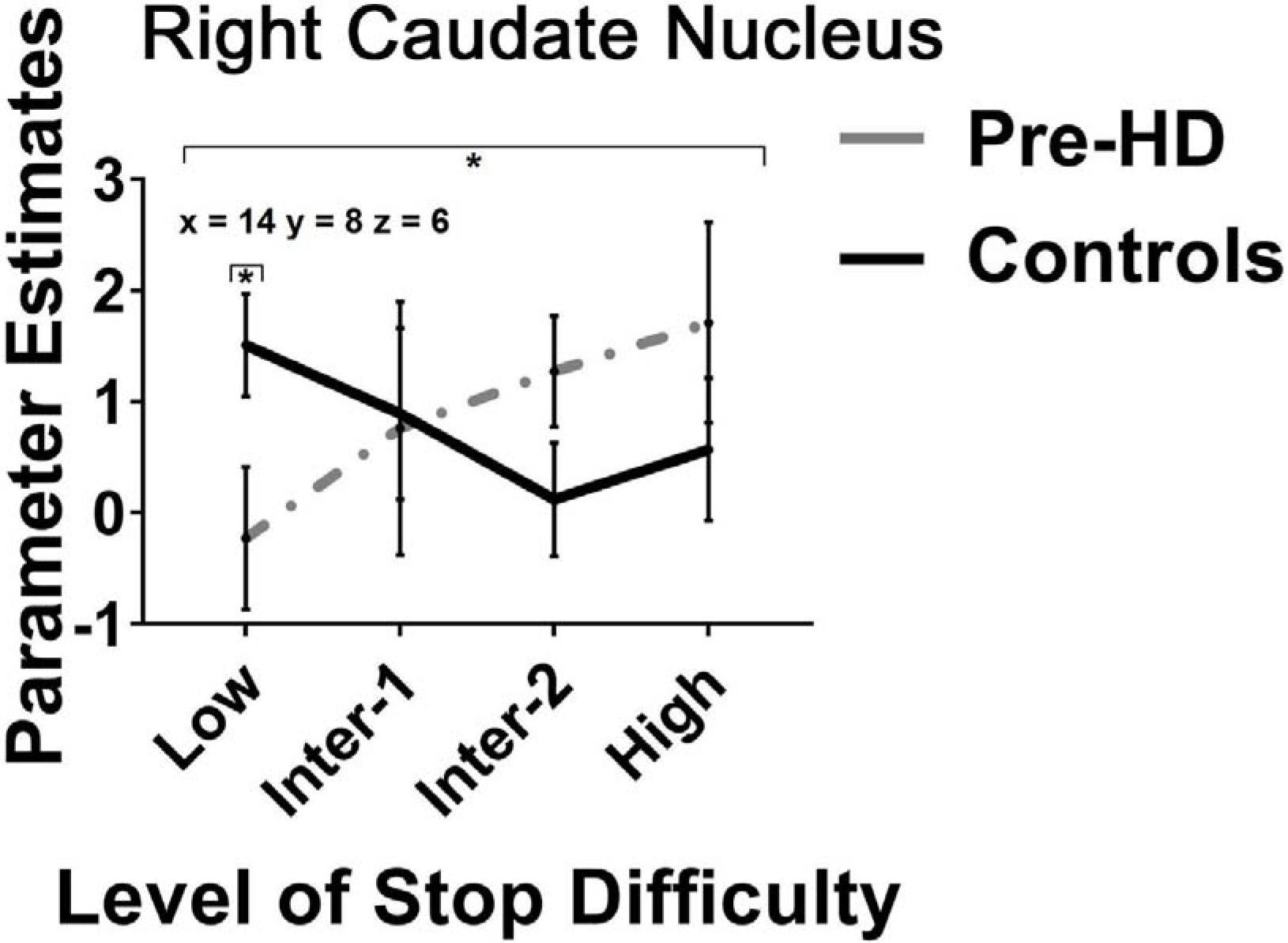
fMRI activity in right caudate nucleus showing significant linear trend for the Group by Level of Stop Difficulty interaction, suggesting that the pattern of fMRI activity in right caudate nucleus varied between pre-HD individuals and controls as a function of stop difficulty. Standard errors are represented by error bars for each group and for each level of stop difficulty. Black asterisks (*) denote a significant linear trend for the Group by Level of Stop Difficulty interaction at *p* < .05.

## 4. Discussion

The primary aim of this study was to test the CRUNCH model (Reuter-Lorenz & Cappell, 2008) to characterise compensation during response inhibition in pre-HD. For this purpose, we modified the standard stop-signal task (Logan & Cowan, 1984) by varying the perceptual load of visual ‘go’ stimuli on ‘go’ trials.

According to the horse-race model, response inhibition can be understood as the outcome of a race between ‘go’ and ‘stop’ processes – if the ‘go’ process finishes before the ‘stop’ process, then response inhibition is unsuccessful; if the ‘stop’ process wins the race, response inhibition is successful (Schmidt et al., 2013; Verbruggen & Logan, 2009). So, if it takes more time for participants to process a perceptually challenging ‘go’ stimulus on ‘go’ trials, then adults will be more likely to successfully inhibit a response on ‘stop’ trials (e.g., Jahfari et al., 2013; Schmidt et al., 2013). So, our modified stop-signal task comprised of four levels of stop difficulty – low perceptual load (high stop difficulty), intermediate-1 perceptual load (intermediate-2 stop difficulty), intermediate-2 perceptual load (intermediate-1 stop difficulty), and high perceptual load (low stop difficulty) – in order to test CRUNCH model predictions (Fabiani, 2012).

The behavioural results showed that, with increased stop difficulty, RTs on ‘go’ trials and the proportion of successful inhibitions on ‘stop’ trials decreased across both groups. These results fit neatly with the notion that response inhibition depends on the relative completion time of ‘go’ and ‘stop’ processes, as predicted by the horse-race model (Verbruggen & Logan, 2009). The results indicate that the modification of the stop-signal paradigm by varying perceptual load of visual ‘go’ stimuli was successful. We confirmed that pre-HD individuals and controls showed comparable ‘go’ RTs, and the pre-HD group had fewer correct responses on ‘go’ trials, compared with controls. Importantly for our aim, we found that pre-HD individuals were significantly slower in inhibiting planned responses, as evidenced by overall longer SSRT, compared with controls. No difference in the number of successful stops (*P*(i)) was observed between pre-HD individuals and controls. Thus, our behavioural results confirm that our task was successful in varying response inhibition difficulty across four levels, and that pre-HD individuals showed largely intact behavioural performance, with some generalised slowing of the stop process.

The CRUNCH model proposes that as task difficulty increases, older adults should show increased fMRI activity compared to younger adults; at higher levels of difficulty (the ‘CRUNCH’ point) the pattern reverses and older adults show less fMRI activity than younger adults. The assumption that underlies this model is that as task difficulty increases, fMRI activity also increases. We did not find this in our task. In contrast with this assumption, controls showed *decreased* fMRI activity with increasing level of stop difficulty in the response inhibition network. Also, contrary to the CRUNCH model, pre-HD individuals showed a linear increase in fMRI activity in right inferior frontal gyrus pars triangularis and right caudate nucleus with increasing stop difficulty, with no evidence of a ‘CRUNCH’ point. No other ROI showed a pattern of fMRI activity that would be consistent with CRUNCH predictions in pre-HD individuals, compared with controls. As such, our results do not support the predictions of the CRUNCH model regarding task difficulty, nor the assumption that fMRI activity increases with task difficulty.

Pre-HD individuals showed overall longer SSRT, compared to controls. Prolonged SSRT in the stop-signal task is usually interpreted as deficient inhibition, and has been reported in Parkinson’s disease (Baglio et al., 2011), obsessive-compulsive disorder (de Wit et al., 2012), schizophrenia (Hughes, Fulham, Johnston, & Michie, 2012), and other psychiatric disorders (Lipszyc & Schachar, 2010). Previous studies also found that pre-HD individuals exhibit deficits in response inhibition on a host of paradigms that require suppression of a response, including the Go/No-Go task (Beste et al., 2011; Bester et al., 2010), the Stroop task (Harrington et al., 2015; Liu et al., 2016; Stout et al., 2011), and the anti-saccade task (Blekher et al., 2006; Golding, Danchaivijitr, Hodgson, Tabrizi, & Kennard, 2006). However, longer SSRT observed in our pre-HD group contrasts with the findings of Rao et al. (2014) who used the stop-signal task and showed no difference in SSRT between pre-HD individuals close and far from predicted disease onset and controls at the behavioural level. Importantly, Rao et al. (2014) did not control for inter-individual variability in the stop-signal delay; however, the stop-signal delay can differentially affect participants’ capacity to inhibit a response on ‘stop’ trials (Hughes et al., 2013; Verbrugenn & Logan, 2008). In contrast, we individually adjusted the starting stop-signal delay for the MRI session on each participants’ response to intermediate-1 level of stop difficulty, so the proportion of successful stops was not affected by variability in the stop-signal delay. Thus, we argue that longer SSRT in our cohort of patients indicate that pre-HD individuals can exhibit deficits in response inhibition *as early as 25 years* prior to predicted disease onset. This novel finding is compatible with functional neuroimaging studies showing that the fronto-striatal circuitry, which governs executive function (Sebastian, Forstmann, & Matzke, 2018) is compromised early in pre-HD (Georgiou-Karistianis et al., 2013a; Poudel et al., 2015; Wolf et al., 2008).

Contrary to the primary assumption of the CRUNCH model, controls did not show increased fMRI activity with increasing stop difficulty in right inferior parietal gyrus pars triangularis and right caudate nucleus. By contrast, controls showed *decreased* fMRI activity with increasing stop difficulty in these brain regions. One reason for the opposite effect to that predicted by the model is perhaps the insensitivity of the CRUNCH model to characterise compensatory processes associated with response inhibition. Importantly, in our recent systematic review of the CRUNCH literature, we (Jamadar, 2018) found that although the CRUNCH is the most influential model of cognitive compensation in the literature, only four studies to date have reported the critical test of the CRUNCH model for fMRI data. We found that these studies investigated the CRUNCH model exclusively in the domain of working memory: verbal working memory (Cappell, Gmeindl, & Reuter-Lorenz, 2010; Schneider-Garces et al., 2010), visuospatial working memory (Bauer, Sammer, & Toepper, 2015; Toepper et al., 2014), and the n-BACK working memory task (Mattay et al., 2006). We concluded that there is evidence for selective reporting of results (the so-called ‘file-drawer effect’) in the CRUNCH literature. Moreover, a recent study that aimed to critically test the CRUNCH model in the domain of visuospatial working memory in pre-HD and controls, showed that as task difficulty increased, fMRI activity also increased (Soloveva et al., 2018). Thus, the failure to meet the primary assumption of the CRUNCH model (that fMRI activity increases with task difficulty) in our study may suggest that the CRUNCH model is not applicable beyond the working memory domain in pre-HD. We suggest that further studies that target various cognitive domains are needed to critically test the CRUNCH model in both health and disease.

Consistent with the argument that CRUNCH does not apply in response inhibition in pre-HD, we also found the opposite effect to that predicted by the CRUNCH model when comparing the pre-HD and control groups. Pre-HD individuals did not show compensatory fMRI overa-ctivation in right inferior frontal gyrus pars triangularis at lower levels of stop difficulty, compared with controls. We also found no evidence of a ‘CRUNCH’ point at higher levels of stop difficulty in pre-HD. These findings are consistent with Rao et al. (2014) who found no difference in the fMRI activity in bilateral inferior frontal gyrus and primary motor cortex during successful inhibition between pre-HD individuals close (< 7.59) and far from (> 12.78) predicted disease onset and controls. Likewise, relative to controls, Unschuld et al. (2012) did not show increased fMRI activity coupled with intact performance on a task that relates to inhibitory control, the Stroop task in pre-HD individuals close (mean of 14.84 years ± 9.65) to predicted disease onset. Right inferior frontal gyrus is a critical node of the ventral attention network, which mediates top-down stimulus-driven control of attention (Corbetta, Patel, & Shulman, 2008). Specifically, the network is involved in re-orienting attention to detect and process behaviorally salient and unexpected stimuli, such as the stop-signal, and facilitate successful stopping (Sebastian et al., 2016; Sharp et al., 2010). The ventral attention network is well preserved in pre-HD (e.g., Hart et al., 2013; Maurage et al., 2017; Wolf et al., 2011) which suggests that inhibitory processes may be therefore less prone to disruption early in the disease. Compatible with this suggestion is the finding that pre-HD individuals and controls did not differ in global grey and white matter volumes in this study. Taken together, it remains possible that the ventral attention network may not be sufficiently impaired in pre-HD individuals to exhibit a pattern of compensatory activity.

However, our behavioural results are not consistent with this conclusion: the finding that pre-HD individuals exhibited significantly longer SSRT relative to controls confirms that our pre-HD group showed impaired response inhibition (e.g., Baglio et al., 2011; Hughes et al., 2012). We suggest that perhaps pre-HD individuals engaged in some compensatory processes to make up for their prolonged SSRT and/or stop difficulty. In favour of this argument, pre-HD individuals showed larger fMRI activity in right inferior frontal gyrus at intermediate-2 and high level of stop difficulty compared to controls, which may be evidence of a compensatory response in pre-HD. Importantly, this pattern of results in pre-HD is consistent with an ‘informal’ definition of neural compensation, and have been often documented in cross-sectional (e.g., Kloppel et al., 2015) and longitudinal studies (e.g., Georgiou-Karistianis et al., 2013a; Poudel et al., 2015).

Our study has several limitations. Firstly, our version of the stop-signal task did not allow us to control for the attentional capture, referred to as the detection of salient signals when inhibition is not yet required (Corbetta et al., 2008; Sharp et al., 2010). Successful stopping in response to an unexpected and salient event, such as stop-signal, requires both that the event is attended and processed, and that the action is inhibited (Sharp et al., 2010). It is therefore important to segregate attentional processes from response inhibition, as this enables to disentangle whether longer SSRT in our cohort of pre-HD individuals, compared with controls, may arise due to deficits in attentional capture rather than due to impaired response inhibition. Furthermore, the pre-HD group was very far from predicted disease onset and showed no significant difference in overall GM and WM volume, compared with controls. Note that while this may have limited compensatory fMRI over-activation at lower levels of stop difficulty, we argue that this does not explain why we did not replicate the CRUNCH effect. Lastly, our sample size is modest (n = 15 pre-HD; n = 15 controls) but is consistent with previous published papers in the pre-HD literature (e.g., n = 15 pre-HD, Kloppel et al., 2009; n = 18 pre-HD, Maurage et al., 2017; n = 16 pre-HD, Wolf et al., 2007; n = 18 pre-HD, Wolf et al., 2011).

This study is the first to show that deficits in response inhibition are observed in pre-HD as early as 25 years prior to predicted disease onset. We conclude that the CRUNCH model does not apply to characterise compensatory processes in pre-HD associated with response inhibition.

## Ethical statement

The study complies with the national ethical research guidelines in Australia. The study was approved by the Monash University and Melbourne Health Human Research Ethics Committees.

## Informed consent

Written informed consent was obtained from each participant in accordance with the Helsinki Declaration.

## Acknowledgement

Jamadar is supported by an Australian Research Council (ARC) Discovery Early Career Research Award (DE150100406) and the ARC Centre of Excellence for Integrative Brain Function (CE140100007).

Poudel was supported by the Hereditary Disease Foundation USA Fellowship and Huntington’s Disease Association (NSW)

We thank all the individuals for participating their time for this study.

## References

Albin, R. L., Reiner, A., Anderson, K. D., Dure, L. S., Handelin, B., Balfour, R., … Young, A. B. (1992). Preferential loss of striato-external pallidal projection neurons in presymptomatic Huntington’s disease. Annals of neurology, 31(4), 425–430.

Andrews, S. C., Domínguez, J. F., Mercieca, E.-C., Georgiou-Karistianis, N., & Stout, J. C. (2015). Cognitive interventions to enhance neural compensation in Huntington’s disease. Neurodegenerative disease management, 5(2), 155–164.

Aron, A. R., Robbins, T. W., & Poldrack, R. A. (2014). Inhibition and the right inferior frontal cortex: one decade on. Trends in Cognitive Sciences, 18(4), 177–185. doi:https://doi.org/10.1016/j.tics.2013.12.003

Ashburner, J., & Friston, K. J. (2001). Why Voxel-Based Morphometry Should Be Used. NeuroImage, 14(6), 1238–1243. doi:https://doi.org/10.1006/nimg.2001.0961

Baglio, F., Blasi, V., Falini, A., Farina, E., Mantovani, F., Olivotto, F., … Bozzali, M. (2011). Functional brain changes in early Parkinson’s disease during motor response and motor inhibition. Neurobiology of Aging, 32(1), 115–124. doi:https://doi.org/10.1016/j.neurobiolaging.2008.12.009

Baltes, P. B., & Lindenberger, U. (1997). Emergence of a powerful connection between sensory and cognitive functions across the adult life span: a new window to the study of cognitive aging? Psychology and aging, 12(1), 12.

Barulli, D., & Stern, Y. (2013). Efficiency, capacity, compensation, maintenance, plasticity: emerging concepts in cognitive reserve. Trends in Cognitive Sciences, 17(10), 502–509.

Bauer, E., Sammer, G., & Toepper, M. (2015). Trying to Put the Puzzle Together: Age and Performance Level Modulate the Neural Response to Increasing Task Load within Left Rostral Prefrontal Cortex. BioMed Research International, 2015, 11. doi:10.1155/2015/415458

Beck, A., Steer, R., & Brown, R. (1996). Manual for Beck Depression Inventory-II.. San Antonio, TX: Psychological Corporation.

Beste, C., Ness, V., Falkenstein, M., & Saft, C. (2011). On the role of fronto-striatal neural synchronization processes for response inhibition—Evidence from ERP phase-synchronization analyses in pre-manifest Huntington’s disease gene mutation carriers. Neuropsychologia, 49(12), 3484–3493. doi:https://doi.org/10.1016/j.neuropsychologia.2011.08.024

Beste, C., Willemssen, R., Saft, C., & Falkenstein, M. (2010). Response inhibition subprocesses and dopaminergic pathways: basal ganglia disease effects. Neuropsychologia, 48(2), 366–373.

Blekher, T., Johnson, S. A., Marshall, J., White, K., Hui, S., Weaver, M., … Foroud, T. (2006). Saccades in presymptomatic and early stages of Huntington disease. Neurology, 67(3), 394.

Brett, M., Anton, J.-L., Valabregue, R., & Poline, J.-B. (2002). Region of Interest Analysis Using an SPM Toolbox [Abstract] (Vol. 16).

Cabeza, R., & Dennis, N. A. (2013). Frontal lobes and aging: deterioration and compensation. In D. T. Stuss & R. T. Knight (Eds.), Principles of Frontal Lobe Function (pp. 628–652). New York: Oxford Univesity Press.

Cappell, K. A., Gmeindl, L., & Reuter-Lorenz, P. A. (2010). Age Differences in Prefontal Recruitment During Verbal Working Memory Maintenance Depend on Memory Load. Cortex; a journal devoted to the study of the nervous system and behavior, 46(4), 462–473. doi:10.1016/j.cortex.2009.11.009

Cieslik, E. C., Mueller, V. I., Eickhoff, C. R., Langner, R., & Eickhoff, S. B. (2015). Three key regions for supervisory attentional control: Evidence from neuroimaging meta-analyses. Neuroscience & Biobehavioral Reviews, 48, 22–34. doi:https://doi.org/10.1016/j.neubiorev.2014.11.003

Corbetta, M., Patel, G., & Shulman, G. L. (2008). The Reorienting System of the Human Brain: From Environment to Theory of Mind. Neuron, 58(3), 306–324. doi:https://doi.org/10.1016/j.neuron.2008.04.017

Donders, F. C. (1969). On the speed of mental processes. Acta Psychologica, 30, 412–431. doi:https://doi.org/10.1016/0001-6918(69)90065-1

Donkers, F. C. L., & van Boxtel, G. J. M. (2004). The N2 in go/no-go tasks reflects conflict monitoring not response inhibition. Brain and Cognition, 56(2), 165–176. doi:https://doi.org/10.1016/j.bandc.2004.04.005

Fabiani, M. (2012). It was the best of times, it was the worst of times: A psychophysiologist’s view of cognitive aging. Psychophysiology, 49(3), 283–304. doi:10.1111/j.1469-8986.2011.01331.x

Fischmeister, F. P. S., Höllinger, I., Klinger, N., Geissler, A., Wurnig, M. C., Matt, E., … Beisteiner, R. (2013). The benefits of skull stripping in the normalization of clinical fMRI data. NeuroImage: Clinical, 3, 369–380. doi:https://doi.org/10.1016/j.nicl.2013.09.007

Friston, K. J., Holmes, A. P., Poline, J. B., Grasby, P. J., Williams, S. C. R., Frackowiak, R. S. J., & Turner, R. (1995). Analysis of fMRI Time-Series Revisited. NeuroImage, 2(1), 45–53. doi:https://doi.org/10.1006/nimg.1995.1007

Georgiou-Karistianis, N. (2009). A peek inside the Huntington’s brain: will functional imaging take us one step closer in solving the puzzle? Experimental Neurology, 220(1), 5–8. doi:https://doi.org/10.1016/j.expneurol.2009.08.001

Georgiou-Karistianis, N., Poudel, G. R., Langmaid, R., Gray, M. A., Churchyard, A., Chua, P., … Stout, J. C. (2013a). Functional and connectivity changes during working memory inHuntington’s disease: 18month longitudinal data from the IMAGE-HD study. Brain and Cognition, 83(1), 80–91.

Georgiou-Karistianis, N., Scahill, R., Tabrizi, S. J., Squitieri, F., & Aylward, E. (2013b). Structural MRI in Huntington’s disease and recommendations for its potential use in clinical trials. Neuroscience & Biobehavioral Reviews, 37(3), 480–490.

Golding, C. V. P., Danchaivijitr, C., Hodgson, T. L., Tabrizi, S. J., & Kennard, C. (2006). Identification of an oculomotor biomarker of preclinical Huntington disease. Neurology, 67(3), 485.

Gray, M., Egan, G., Ando, A., Churchyard, A., Chua, P., Stout, J., & Georgiou-Karistianis, N. (2013). Prefrontal activity in Huntington’s disease reflects cognitive and neuropsychiatric disturbances: the IMAGE-HD study. Experimental Neurology, 239, 218–228.

Gregory, S., Long, J. D., Klöppel, S., Razi, A., Scheller, E., Minkova, L., … Track-On, i. (2018). Testing a longitudinal compensation model in premanifest Huntington’s disease. Brain, awy122–awy122. doi:10.1093/brain/awy122

Gregory, S., Long, J. D., Klöppel, S., Razi, A., Scheller, E., Minkova, L., … Leavitt, B. R. (2017). Operationalizing compensation over time in neurodegenerative disease. Brain, 140(4), 1158–1165.

Harrington, D. L., Rubinov, M., Durgerian, S., Mourany, L., Reece, C., Koenig, K., … Rao, S. M. (2015). Network topology and functional connectivity disturbances precede the onset of Huntington’s disease. Brain, 138(8), 2332–2346. doi:10.1093/brain/awv145

Hart, E. P., Dumas, E. M., Zwet, E. W., Hiele, K., Jurgens, C. K., Middelkoop, H. A. M., … Roos, R. A. C. (2013). Longitudinal pilot-study of Sustained Attention to Response Task and P300 in manifest and pre-manifest Huntington’s disease. Journal of Neuropsychology, 9(1), 10–20. doi:10.1111/jnp.12031

Hua, J., Unschuld, P. G., Margolis, R. L., Zijl, P., & Ross, C. A. (2014). Elevated arteriolar cerebral blood volume in prodromal Huntington’s disease. Movement Disorders, 29(3), 396–401.

Hughes, M. E., Fulham, W. R., Johnston, P. J., & Michie, P. T. (2012). Stop-signal response inhibition in schizophrenia: Behavioural, event-related potential and functional neuroimaging data. Biological Psychology, 89(1), 220–231. doi:https://doi.org/10.1016/j.biopsycho.2011.10.013

Hughes, M. E., Johnston, P. J., Fulham, W. R., Budd, T. W., & Michie, P. T. (2013). Stop-signal task difficulty and the right inferior frontal gyrus. Behavioural Brain Research, 256, 205–213. doi:https://doi.org/10.1016/j.bbr.2013.08.026

Iannetti, G. D., & Wise, R. G. (2007). BOLD functional MRI in disease and pharmacological studies: room for improvement? Magnetic Resonance Imaging, 25(6), 978–988. doi:https://doi.org/10.1016/j.mri.2007.03.018

Jahfari, S., Ridderinkhof, K. R., & Scholte, H. S. (2013). Spatial Frequency Information Modulates Response Inhibition and Decision-Making Processes. PLOS ONE, 8(10), e76467. doi:10.1371/journal.pone.0076467

Ji, L., Pearlson, G. D., Hawkins, K. A., Steffens, D. C., Guo, H., & Wang, L. (2018). A New Measure for Neural Compensation Is Positively Correlated With Working Memory and Gait Speed. Frontiers in Aging Neuroscience, 10(71). doi:10.3389/fnagi.2018.00071

Kloppel, S., Draganski, B., Siebner, H. R., Tabrizi, S. J., Weiller, C., & Frackowiak, R. S. (2009). Functional compensation of motor function in pre-symptomatic Huntington’s disease. Brain, 132(Pt 6), 1624–1632. doi:10.1093/brain/awp081

Kloppel, S., Gregory, S., Scheller, E., Minkova, L., Razi, A., Durr, A., … Landwehrmeyer, G. B. (2015). Compensation in preclinical Huntington’s disease: evidence from the track-on HD study. EBioMedicine, 2(10), 1420–1429.

Langbehn, D. R., Brinkman, R. R., Falush, D., Paulsen, J. S., & Hayden, M. R. (2004). A new model for prediction of the age of onset and penetrance for Huntington’s disease based on CAG length. Clinical Genetics, 65(4), 267–277. doi:10.1111/j.1399-0004.2004.00241.x

Levy, B. J., & Wagner, A. D. (2011). Cognitive control and right ventrolateral prefrontal cortex: reflexive reorienting, motor inhibition, and action updating. Annals of the New York Academy of Sciences, 1224(1), 40–62. doi:10.1111/j.1749-6632.2011.05958.x

Lipszyc, J., & Schachar, R. (2010). Inhibitory control and psychopathology: A meta-analysis of studies using the stop signal task. Journal of the International Neuropsychological Society, 16(6), 1064–1076. doi:10.1017/S1355617710000895

Liu, W., Yang, J., Chen, K., Luo, C., Burgunder, J., Gong, Q., & Shang, H. (2016). Resting-state fMRI reveals potential neural correlates of impaired cognition in Huntington’s disease. Parkinsonism & Related Disorders, 27, 41–46. doi:https://doi.org/10.1016/j.parkreldis.2016.04.017

Logan, G. D., & Cowan, W. B. (1984). On the ability to inhibit thought and action: A theory of an act of control. Psychological Review, 91(3), 295–327. doi:10.1037/0033-295X.91.3.295

MacDonald, Ambrose, C. M., Duyao, M. P., Myers, R. H., Lin, C., Srinidhi, L., … Groot, N. (1993). A novel gene containing a trinucleotide repeat that is expanded and unstable on Huntington’s disease chromosomes. Cell, 72(6), 971–983.

Malejko, K., Weydt, P., Süßmuth, S. D., Grön, G., Landwehrmeyer, B. G., & Abler, B. (2014). Prodromal Huntington Disease as a Model for Functional Compensation of Early Neurodegeneration. PLOS ONE, 9(12), e114569. doi:10.1371/journal.pone.0114569

Mattay, V. S., Fera, F., Tessitore, A., Hariri, A. R., Berman, K. F., Das, S., … Weinberger, D. R. (2006). Neurophysiological correlates of age-related changes in working memory capacity. Neuroscience Letters, 392(1), 32–37. doi:https://doi.org/10.1016/j.neulet.2005.09.025

Maurage, P., Heeren, A., Lahaye, M., Jeanjean, A., Guettat, L., Verellen-Dumoulin, C., … Constant, E. (2017). Attentional Impairments in Huntington’s Disease: A Specific Deficit for the Executive Conflict (Vol. 31).

Nelson, H., Willison, T., & Owen Adrian, M. (1992). National adult reading test (2nd edition). International Journal of Geriatric Psychiatry, 7(7), 533.

Orth, M., & Schwenke, C. (2011). Age-at-onset in Huntington disease. PLoS Currents, 3, RRN1258. doi:10.1371/currents.RRN1258

Poudel, G., Stout, J. C., Gray, M. A., Salmon, L., Churchyard, A., Chua, P., … Georgiou-Karistianis, N. (2015). Functional changes during working memory in Huntington’s disease: 30-month longitudinal data from the IMAGE-HD study. Brain Structure and Function, 220(1), 501–512.

Rao, J. A., Harrington, D. L., Durgerian, S., Reece, C., Mourany, L., Koenig, K., … Rao, S. M. (2014). Disruption of response inhibition circuits in prodromal Huntington disease. Cortex; a journal devoted to the study of the nervous system and behavior, 58, 72–85. doi:10.1016/j.cortex.2014.04.018

Reid, L. B., Boyd, R. N., Cunnington, R., & Rose, S. E. (2016). Interpreting Intervention Induced Neuroplasticity with fMRI: The Case for Multimodal Imaging Strategies. Neural Plasticity, 2016, 13. doi:10.1155/2016/2643491

Reuter-Lorenz, P. A., & Cappell, K. A. (2008). Neurocognitive aging and the compensation hypothesis. Current directions in psychological science, 17(3), 177–182.

Rosas, Salat, D. H., Lee, S. Y., Zaleta, A. K., Pappu, V., Fischl, B., … Hersch, S. M. (2008). Cerebral cortex and the clinical expression of Huntington’s disease: complexity and heterogeneity. Brain, 131(4), 1057–1068.

Scheller, E., Minkova, L., Leitner, M., & Klöppel, S. (2014). Attempted and successful compensation in preclinical and early manifest neurodegeneration–a review of task fMRI studies. Frontiers in psychiatry, 5.

Schmidt, R., Leventhal, D. K., Mallet, N., Chen, F., & Berke, J. D. (2013). Canceling actions involves a race between basal ganglia pathways. Nature Neuroscience, 16, 1118. doi:10.1038/nn.3456 https://www.nature.com/articles/nn.3456#supplementary-information

Schneider-Garces, N. J., Gordon, B. A., Brumback-Peltz, C. R., Shin, E., Lee, Y., Sutton, B. P., … Fabiani, M. (2010). Span, CRUNCH, and Beyond: Working Memory Capacity and the Aging Brain. Journal of cognitive neuroscience, 22(4), 655–669. doi:10.1162/jocn.2009.21230

Sebastian, A., Forstmann, B. U., & Matzke, D. (2018). Towards a model-based cognitive neuroscience of stopping – a neuroimaging perspective. Neuroscience & Biobehavioral Reviews, 90, 130–136. doi:https://doi.org/10.1016/j.neubiorev.2018.04.011

Sebastian, A., Jung, P., Neuhoff, J., Wibral, M., Fox, P. T., Lieb, K., … Mobascher, A. (2016). Dissociable attentional and inhibitory networks of dorsal and ventral areas of the right inferior frontal cortex: a combined task-specific and coordinate-based meta-analytic fMRI study. Brain Structure and Function, 221(3), 1635–1651. doi:10.1007/s00429-015-0994-y

Sharp, D. J., Bonnelle, V., De Boissezon, X., Beckmann, C. F., James, S. G., Patel, M. C., & Mehta, M. A. (2010). Distinct frontal systems for response inhibition, attentional capture, and error processing. Proceedings of the National Academy of Sciences, 107(13), 6106.

Smith, H. (1991). Symbol Digit Modalities Test. Los Angeles: Western Psychological Services.

Smith, J. L., Jamadar, S., Provost, A. L., & Michie, P. T. (2013). Motor and non-motor inhibition in the Go/NoGo task: An ERP and fMRI study. International Journal of Psychophysiology, 87(3), 244–253. doi:https://doi.org/10.1016/j.ijpsycho.2012.07.185

Smith, J. L., Johnstone, S. J., & Barry, R. J. (2007). Response priming in the Go/NoGo task: The N2 reflects neither inhibition nor conflict. Clinical Neurophysiology, 118(2), 343–355. doi:https://doi.org/10.1016/j.clinph.2006.09.027

Soloveva, M. V., Jamadar, S. D., Poudel, G., & Georgiou-Karistianis, N. (2018). A critical review of brain and cognitive reserve in Huntington’s disease. Neuroscience & Biobehavioral Reviews, 88, 155–169. doi:https://doi.org/10.1016/j.neubiorev.2018.03.003

Soloveva, M. V., Jamadar, S. D., Velakoulis, D., Poudel, G., & Georgiou-Karistianis, N. (2018). To CRUNCH or not to CRUNCH: Task Difficulty Affects Functional Brain Reorganisation during Visuospatial Working Memory Performance in Premanifest Huntington’s Disease. bioRxiv. doi:10.1101/459180

Steffener, J., & Stern, Y. (2012). Exploring the neural basis of cognitive reserve in aging. Biochimica et Biophysica Acta (BBA)-Molecular Basis of Disease, 1822(3), 467–473.

Stella J. de Wit, Froukje E. de Vries, Ysbrand D. van der Werf, Danielle C. Cath, Dirk J. Heslenfeld, Eveline M. Veltman, … Odile A. van den Heuvel. (2012). Presupplementary Motor Area Hyperactivity During Response Inhibition: A Candidate Endophenotype of Obsessive-Compulsive Disorder. American Journal of Psychiatry, 169(10), 1100–1108. doi:10.1176/appi.ajp.2012.12010073

Stout, J. C., Paulsen, J. S., Queller, S., Solomon, A. C., Whitlock, K. B., Campbell, J. C., … Aylward, E. H. (2011). Neurocognitive signs in prodromal Huntington disease. Neuropsychology, 25(1), 1–14. doi:10.1037/a0020937

Tabrizi, S. J., Langbehn, D. R., Leavitt, B. R., Roos, R. A., Durr, A., Craufurd, D., … Scahill, R. I. (2009). Biological and clinical manifestations of Huntington’s disease in the longitudinal TRACK-HD study: cross-sectional analysis of baseline data. The Lancet Neurology, 8(9), 791–801.

Toepper, M., Gebhardt, H., Bauer, E., Haberkamp, A., Beblo, T., Gallhofer, B., … Sammer, G. (2014). The impact of age on load-related dorsolateral prefrontal cortex activation. Frontiers in Aging Neuroscience, 6(9). doi:10.3389/fnagi.2014.00009

Verbruggen, F., & Logan, G. D. (2008). Response inhibition in the stop-signal paradigm. Trends in Cognitive Sciences, 12(11), 418–424. doi:https://doi.org/10.1016/j.tics.2008.07.005

Verbruggen, F., & Logan, G. D. (2009). Models of response inhibition in the stop-signal and stop-change paradigms. Neuroscience & Biobehavioral Reviews, 33(5), 647–661. doi:https://doi.org/10.1016/j.neubiorev.2008.08.014

Wolf, R. C., Grön, G., Sambataro, F., Vasic, N., Wolf, N. D., Thomann, P. A., … Orth, M. (2011). Brain activation and functional connectivity in premanifest Huntington’s disease during states of intrinsic and phasic alertness. Human Brain Mapping, 33(9), 2161–2173. doi:10.1002/hbm.21348

Wolf, R. C., Sambataro, F., Vasic, N., Schönfeldt-Lecuona, C., Ecker, D., & Landwehrmeyer, B. (2008). Altered frontostriatal coupling in pre-manifest Huntington’s disease: effects of increasing cognitive load. European Journal of Neurology, 15(11), 1180–1190. doi:10.1111/j.1468-1331.2008.02253.x

